# Bola-amphiphilic dendrimer empowers imatinib to target metastatic ovarian cancer stem cells *via* β-catenin-HRP2 signaling axis

**DOI:** 10.1101/2024.06.07.597857

**Authors:** Zeyu Shi, Margarita Artemenko, Ming Zhang, Canhui Yi, Peng Chen, Shuting Lin, Zhancun Bian, Baoping Lian, Fanzhen Meng, Jiaxuan Chen, Tom Roussel, Ying Li, Karen K. L. Chan, Philip P. C. Ip, Hung-Cheng Lai, Xiaoxuan Liu, Ling Peng, Alice S. T. Wong

## Abstract

Ovarian cancer is the leading cause of death among all gynecological malignancies, and drug resistance renders the current chemotherapy agents ineffective for patients with advanced metastatic tumors. We report an effective treatment strategy for targeting metastatic ovarian cancer involving a nanoformulation (Bola/IM) – bola-amphiphilic dendrimer (Bola)-encapsulated imatinib (IM) – to target the critical mediator of ovarian cancer stem cells (CSCs) CD117 (c-Kit). Bola/IM offered significantly more effective targeting of CSCs compared to IM alone, through a novel and tumor-specific β-catenin/HRP2 axis, allowing potent inhibition of cancer cell survival, stemness and metastasis in metastatic and drug-resistant ovarian cancer cells. Promising results were also obtained in clinically relevant patient-derived ascites and organoids, alongside high tumor-oriented accumulation and favorable pharmacokinetic properties in mouse models. Furthermore, Bola/IM displayed synergistic anticancer activity when combined with the first-line chemotherapeutic drug cisplatin in patient-derived xenograft mouse models, without any adverse effects. Our findings support the use of Bola/IM as a nanoformulation to empower IM, providing targeted and potent treatment of metastatic ovarian cancer. Our study thus represents a significant advancement towards addressing the unmet medical need for improved therapies targeting this challenging disease.

## INTRODUCTION

Ovarian cancer is the most lethal gynecological malignancy worldwide [1, 2] . Being asymptomatic, more than 75% of ovarian cancer cases are diagnosed at an advanced metastatic stage [3]. Platinum/taxane-based chemotherapy is the first-line standard approach used for treating unresectable ovarian cancer [4] but is greatly hindered by intrinsic and acquired chemoresistance. While metastatic tumors are initially sensitive to this chemotherapy, 70% of patients will experience recurrence, which is the major cause of mortality, with a 5-year survival rate of <25% [5]. Thus, there is an urgent need for new therapeutic strategies that can provide increased specificity and efficacy in treating metastatic, drug-resistant ovarian cancer.

Cancer stem cells (CSCs) are a small subpopulation of tumor cells that possess a high capacity for self-renewal and multilineage differentiation, tumor initiation, chemoresistance, and metastasis [6]. As such, targeting CSCs represents a critically important strategy in treating metastatic, chemoresistant cancers. We have reported a critical role for the receptor tyrosine kinase c-Kit (also known as CD117) in CSCs. Not only does c-Kit serve as a marker of ovarian CSCs, but it also determines their stem phenotype [7]. While activating c-Kit mutations have not been found in ovarian cancer, increased c-Kit expression has been correlated with poorer patient outcomes [7]. Moreover, c-Kit is hyperactivated in tumor-infiltrating immune cells [8], suggesting that targeting c-Kit may also relieve immunosuppression in the tumor microenvironment.

One ideal candidate for targeting c-Kit is imatinib (IM), an FDA-approved anticancer drug shown to inhibit c-Kit, Bcr/Abl, and platelet-derived growth factor receptor [9]. However, clinical trials have shown that IM alone or in combination with docetaxel/paclitaxel has little or no positive effect in overall response or progression free survival rates in patients with primary or recurrent ovarian cancer (www.clinicaltrials.gov: NCT00510653, NCT00216112, NCT00840450). Furthermore, while the use of a daily dose of 400 mg IM is generally well-tolerated, there has been rare but nevertheless lethal hematological and hepato/cardiac toxicities that also need to be considered and addressed [10]. A lower dose regimen allowing safe and effective antitumor efficacy is therefore urgently needed.

In recent years, there has been significant progress in the development of nanotechnology-based drug delivery to enhance the effectiveness of anticancer drugs while minimizing their side effects [11–13] by passively targeting the tumor *via* the enhanced permeability and retention (EPR) effect [14–16]. This phenomenon is unique to the tumor microenvironment in which a leaky vasculature and dysfunctional lymphatic drainage promotes the accumulation and retention of nanosized particles within the tumor lesion. A myriad of nanomaterials such as lipid nanoparticles, polymers, peptides, and proteins have thus been developed to exploit this EPR effect for tumor-targeted drug delivery in cancer treatment [11,13].

Dendrimers, a special class of synthetic polymers with radially symmetric and well-defined structures, have emerged as high precision materials in drug delivery by virtue of their precisely controllable structure and cooperative multivalence [17, 18]. In particular, amphiphilic dendrimers have proven to be very useful in the construction of modular supramolecular nanosystems for drug delivery [19, 20]. Composed of distinct hydrophobic chains and hydrophilic dendrons, these amphiphilic dendrimers are able to self-assemble into nanoassemblies similar to lipids while retaining their unique dendrimer structural properties [21–24]. We previously developed bola-amphiphilic poly(amidoamine) (PAMAM) dendrimers for nucleic acid delivery, comprising a hydrophobic “bola-lipid” core and two hydrophilic dendron terminals [25, 26]. We designed the bola-amphiphilic scaffold to mimic the strong assembly properties of bola-amphiphiles found in extremophile archae bacteria [27]. To enable cancer cell-specific delivery, we incorporated a thioacetal group into our bola-amphiphilic dendrimers thus rendering them responsive to the high level of reactive oxygen species (ROS) in cancer cells [25, 26]. Fluorine tags introduced at the core of the dendrimer scaffold also enables the tracking of ROS-responsive delivery using ^19^F-NMR [26, 28].

In this study, we report for the first time the ability of the bola-amphiphilic dendrimer of generation 2 (referred to hereafter as Bola) to encapsulate the anticancer drug IM and form a stable nanoparticulate formulation (Bola/IM) (Scheme 1). Importantly, low doses of Bola/IM, *via* a novel β-catenin/HRP2 axis, more effectively targeted ovarian CSCs compared to high dose IM treatment alone and avoided toxicity in a mouse model of metastatic ovarian cancer. In addition, Bola/IM combined with the chemotherapeutic drug cisplatin demonstrated a synergistic anticancer effect in patient-derived ascites models, with no adverse side effects. These results highlight the therapeutic potential of Bola/IM in metastatic ovarian cancer for which an efficacious treatment is urgently required.

**Scheme 1.**
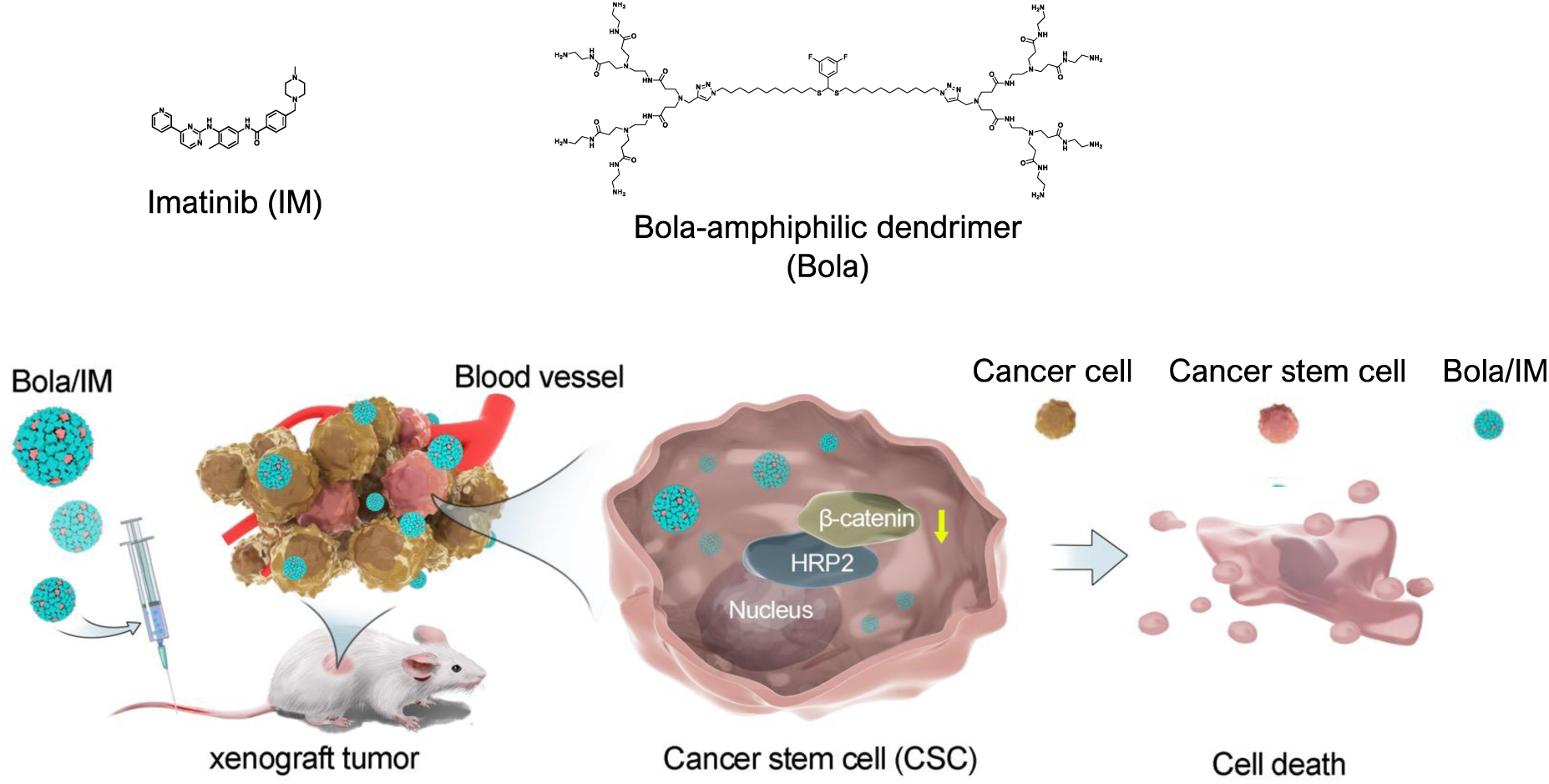

## RESULTS

### Robust Nanoformulation of IM with the Bola-Amphiphilic Dendrimer (Bola)

We have successfully exploited amphiphilic dendrimers for the encapsulation of structurally diverse anticancer drugs including doxorubicin, paclitaxel and rapamycin previously. These dendrimers self-assemble into supramolecular dendrimer nanomicelles with spacious interior cavities for excellent high drug-loading and encapsulation efficiencies [22–24]. The dendrimers used in these earlier studies were classical amphiphiles composed of a hydrophobic chain component and a hydrophilic PAMAM dendron part. In the present study, we investigated a novel approach employing bola-amphiphilic dendrimers of generation 2 (Bola) [25,26] for drug encapsulation. Unlike the classical amphiphile dendrimers, Bola features a unique structure comprising a bola-lipid thioacetal core and two PAMAM dendrons at each of two terminals. This distinctive composition holds the potential to form stable formulations for drug encapsulation while allowing ROS-responsive drug release, offering exciting possibilities for further exploration. Previous studies using ^1^H/^13^C/^19^F-NMR, high-resolution mass spectrometry and HPLC analysis have permitted the full characterization of Bola.

We first studied the drug loading capacity and encapsulation efficiency of IM by Bola using various ratios of IM and Bola (w/w) (16/2, 16/4 and 16/8). Specifically, the thin film dispersion method was used to encapsulate IM by Bola [22]. Our findings revealed a maximum drug-loading efficiency of 33%, and drug encapsulation efficiency ranging from 96% to 100% (Figure S1a). These results highlight that Bola effectively encapsulates IM with high loading and encapsulation capacities.

The IM-loaded dendrimer assemblies (Bola/IM) obtained in this study exhibited small size and spherical shape, as revealed using transmission electron microscopy (TEM) (Figure 1a). Dynamic light scattering (DLS) analysis further confirmed the small size of the Bola/IM, with an average diameter of approximately 30 nm (PDI: ∼0.2) (Figure S1b) and a zeta-potential of +33 (±3) mV (Figure S1c). These findings collectively demonstrate the successful Bola-induced nanoformulation of IM, emphasizing its effectiveness in producing small, stable and well-defined nanoparticles.

**Figure 1.**
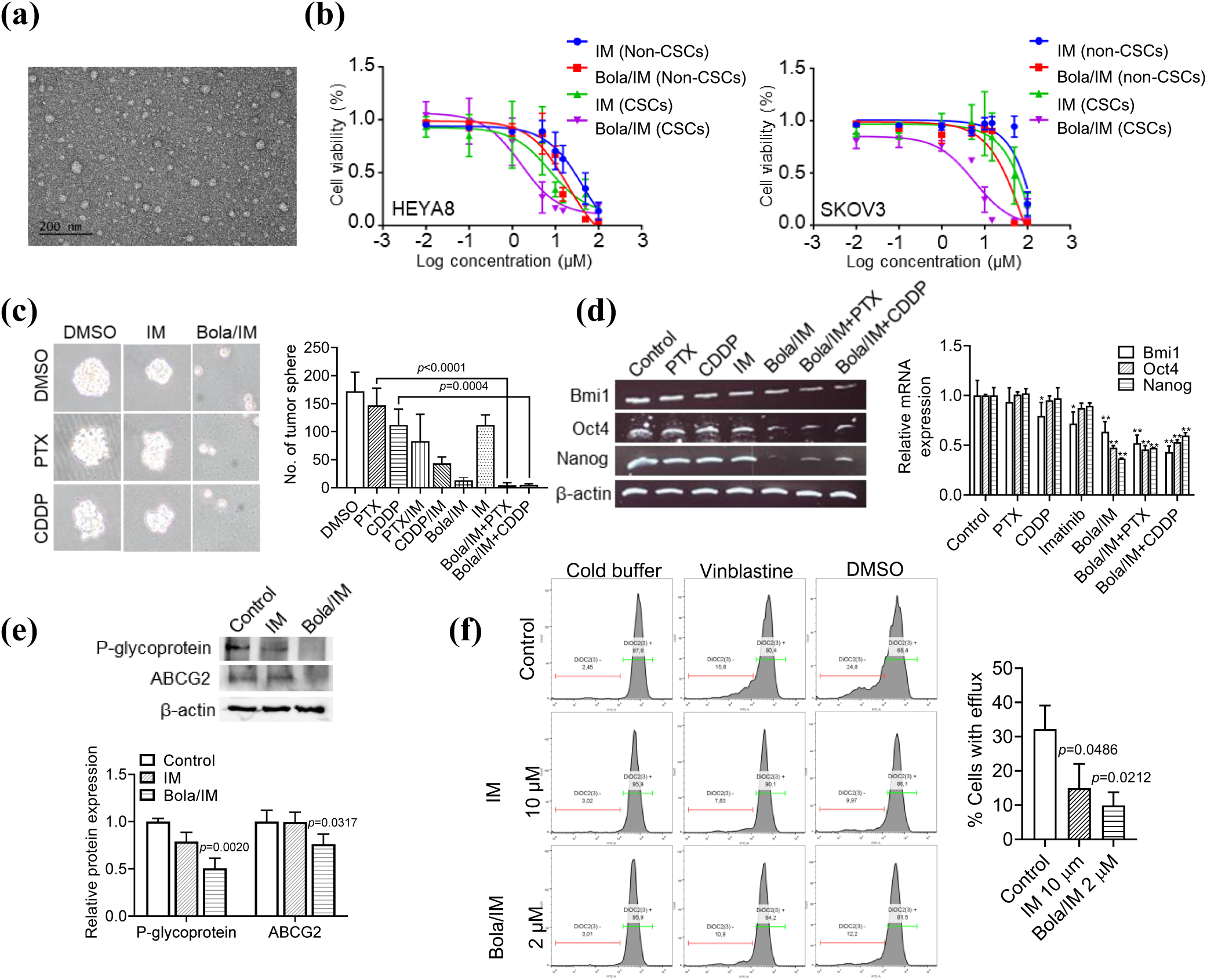
Bola/IM nanoformulation inhibits CSCs *in vitro*. (a) TEM images of Bola/IM. (b) *In vitro* cytotoxicity IC_50_ (μM) values of IM, Bola and Bola/IM towards non-CSCs and CSCs. (c) Spheroids treated with paclitaxel (PTX) or cisplatin (CDDP) with/without IM or Bola/IM. Pictures were taken at Day 7. Number of tumor spheres were counted. (d) Changes of stemness markers were identified by RT-PCR in HEYA8 CSCs. β-actin was used as a control. (e) Cells treated with IM or Bola/IM and expression of P-glycoprotein and ABCG2 were detected by Western blot. β-actin was used as a control. (f) Bola/IM nanoformulation reduced drug efflux in HEYA8 CSCs. Experiments were repeated three times and data are shown as mean ± SD. *, *P*<0.05 vs control. ***, *P*<0.001 vs control.

### Bola/IM Nanoformulation Inhibits CSC *In Vitro*

With the stable and robust Bola/IM nanoformulation in hand, we first conducted MTT assays to evaluate its therapeutic potential in targeting both CSCs and non-CSCs *in vitro*. Strikingly, the results demonstrated that compared to non-CSCs, CSCs exhibited up to 16-fold higher sensitivity to Bola/IM (Figure 1b; Figure S1d). The IC_50_ for HEYA8 CSCs and SKOV3 CSCs were determined to be 1.6 μM and 5.7 μM, respectively; in contrast to 19 μM for HEYA8 non-CSCs and 96 μM for SKOV3 non-CSCs, and 39 μM for normal ovarian surface epithelial (OSE) cells. The significant difference in effective doses between OSE and CSCs suggests that Bola/IM exhibits favorable tolerance in eliminating CSCs. Compared to IM, Bola/IM demonstrated superior efficacy compared to IM for both CSCs and non-CSCs, yet with remarkably higher potency to target CSCs over non-CSCs (Figure 1b; Figure S1d).

To further investigate the therapeutic efficacy of Bola/IM in treating CSCs, we employed sphere formation assays, which are a functional hallmark of CSC renewal. The results indicated that Bola/IM was significantly more effective in blocking CSC self-renewal compared to naked IM at the same dose (Figure 1c). Moreover, Bola/IM sensitized chemoresistant CSCs to paclitaxel (PTX) and cisplatin (CDDP) (Figure 1c). Notably, treatment with Bola/IM led to marked decreases in CSC stemness markers such as Bmi1, Nanog and Oct4 in comparison to treatment with IM, PTX or CDDP alone (Figure 1d). In contrast, naked Bola had no effect on cell viability or CSC renewal ability (Figure S2a, b, c).

Given the potential association between CSC-driven chemoresistance and overexpression of drug efflux pumps on the cell membrane [29], we investigated the impact of Bola/IM on the two major efflux pumps in ovarian CSCs, ABCG2 and P-glycoprotein. Remarkably, treatment with Bola/IM resulted in a considerable decrease in the levels of ABCG2 and P-glycoprotein, whereas equivalent concentrations of IM exhibited no such effect (Figure 1e). These findings suggest that Bola/IM effectively inhibits drug efflux, which could explain its ability to overcome chemoresistance (Figure 1f). In contrast, naked Bola demonstrated no reduction in drug efflux (Figure S2d). These findings demonstrating the ability of Bola/IM to interfere with CSC self-renewal and overcome chemoresistance by inhibiting drug efflux pumps altogether highlight Bola/IM as a promising effective therapeutic strategy for targeting CSCs.

### Bola/IM Inhibits the β-catenin/HRP2 Signaling Pathway

We previously reported that c-Kit could activate β-catenin signaling in mediating chemoresistance in CSCs [7]. Given the effectiveness of Bola/IM in treating CSCs but not non-CSCs while overcoming drug resistance, we sought to investigate whether Bola/IM was involved in the c-Kit activated β-catenin signaling pathway. To compare the differential interacting partners of β-catenin downstream of c-Kit between non-CSCs and CSCs, we performed co-immunoprecipitation (co-IP) followed by LC-MS/MS proteome analysis.

Among the potential partners identified, the hepatoma-derived growth factor related protein 2 (HRP2) was found to be among the most significantly enriched proteins in CSCs compared to non-CSCs (9.84-fold). The interaction between β-catenin and HRP2 was confirmed by co-IP and western blot analysis (Figure 2a). The differential levels of β-catenin-HRP2 interaction levels between CSCs and non-CSCs may be attributed to the higher basal expression of HRP2 in CSCs (Figure 2b). HRP2 expression was lost upon CSC differentiation and was significantly lower in normal ovarian surface epithelium (OSE) (Figure 2b). Immunofluorescence microscopy further confirmed the nuclear localization of HRP2 (Figure 2c).

**Figure 2.**
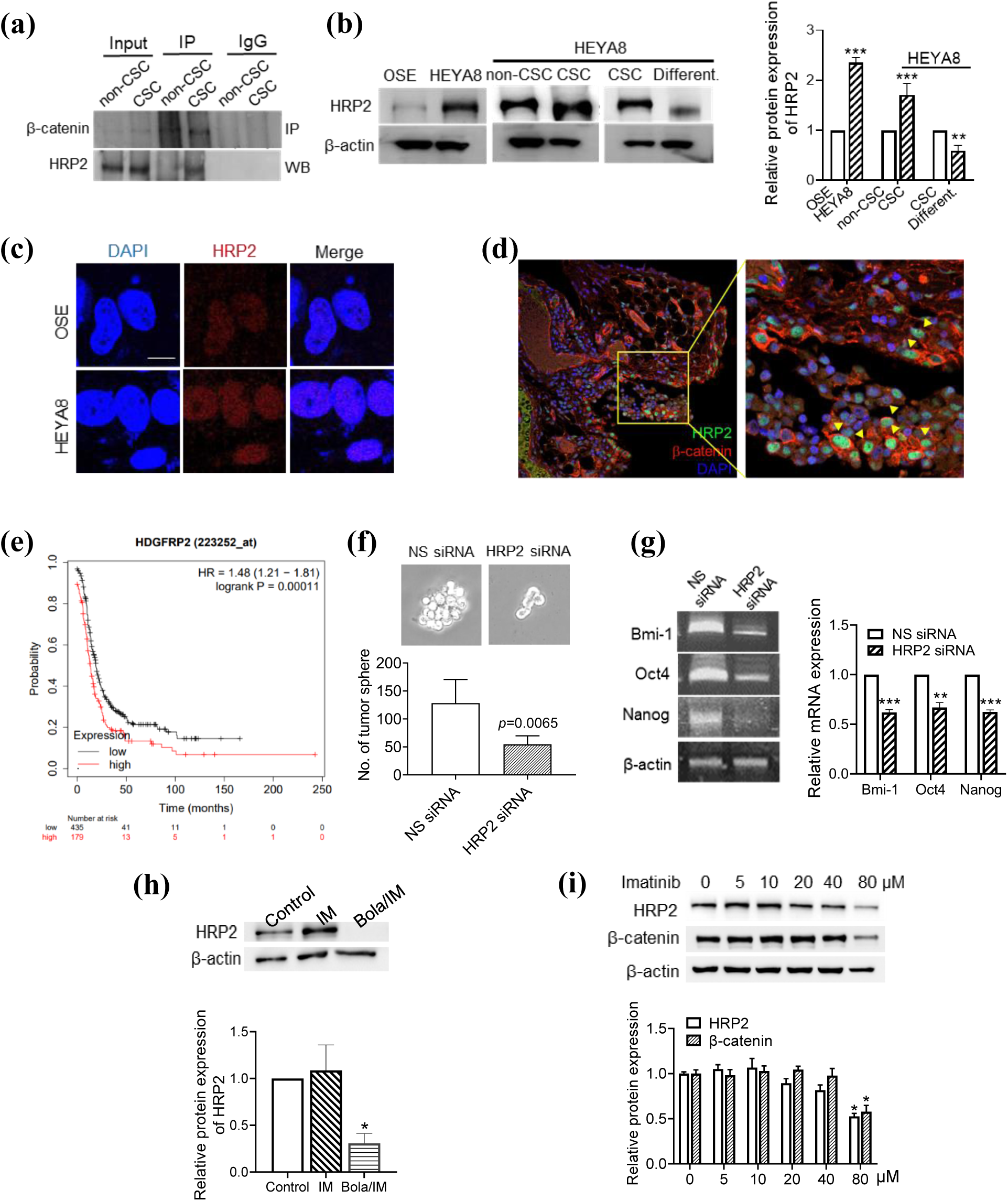
Bola/IM inhibits the β-catenin/HRP2 signaling pathway. (a) HRP2/β-catenin complexes were detected by Western blot using anti-HRP2 antibody after immunoprecipitation by β-catenin primary antibody with nuclear extracts. (b) Western blot analysis of HRP2 expression in OSE, HEYA8, and HEYA8 CSCs, and differentiated cells from spheroids. β-actin was used as a control. (c) The expression and distribution of HRP2 in OSE and HEYA8 through immunofluorescence analysis. (d) Nuclear localization of HRP2 and β-catenin in patient sample were visualized by immunofluorescence on cryosection (yellow arrows). (e) Kaplan-Meier analysis of progression-free survival with high or low HRP2 expression for ovarian cancer patients (*P*=0.00011, log-rank test). (f) Spheroids were treated with 20 nM HRP2 siRNA or control siRNA and the representative images of tumor spheroids were recorded. (g) Changes in stemness markers were identified by RT-PCR. β-actin was used as a control. (h) Western blot analysis of HRP2 expression upon IM or Bola/IM treatment a1**t**25 μM. (i) Western blot analysis of HRP2 and β-catenin expression upon IM treatment at dose series. β-actin was used as a control. Experiments were repeated three times and data are shown as mean ± SD. *, *P*<0.05 vs control. **, *P*<0.01 vs control. ***, *P*<0.001 vs control.

Also importantly, we observed a positive association between HRP2 and β-catenin in clinical samples (Figure 2d). High expression of HRP2 also tended to correlate with poor clinical outcomes (Figure 2e). In addition, HRP2 siRNA suppressed CSC sphere formation (Figure 2f), and the expression of stemness markers (Bmi-1, Oct4, and Nanog) (Figure 2g). Notably, treatment with Bola/IM at 5 μM significantly inhibited HRP2 expression (Figure 2h), whereas a similar decrease in HRP2 expression was only observed with naked IM at very high concentration (80 μM) (Figure 2i). These findings are consistent with our previous observations regarding the effect of c-Kit on CSC growth inhibition and provide further evidence of the significantly more pronounced effect of Bola/IM on CSCs compared to that of naked IM. The data also highlight the potential of Bola/IM in targeting the c-Kit/β-catenin-HRP2 signaling pathway in CSCs, which could contribute to its superior therapeutic efficacy.

### Therapeutic Effects of Bola/IM Nanoformulation *In Vivo*

To evaluate the anticancer activity of Bola/IM *in vivo*, we used a mouse model of metastatic ovarian cancer induced by CSCs (Figure 3a). HEYA8 CSCs were injected intra-peritoneally into mice to generate the metastatic ovarian cancer model, and Bola/IM, or PBS or IM alone as control, then administrated intravenously. Remarkably, even at a low dose of 1.0 mg/kg, Bola/IM significantly suppressed peritoneal metastases (Figure 3b), while IM alone only showed inhibition at the 5-fold higher dose of 5.0 mg/kg but not at the 2.5-fold higher dose of 2.5 mg/kg (Figure 3b; Figure S3a), suggesting increased effectivenesss of Bola/IM compared to IM alone. Furthermore, the number of tumor nodules was significantly reduced after treatment with Bola/IM (Figure 3c). Importantly, Bola/IM was well tolerated by the mice, as we observed no abnormal behavior or weight loss during the entire treatment period (Figure S3b; Figure S4). In addition, we found no pathological alterations in blood, bone marrow, or peritoneal fluid (Figure 3d). Also, histopathological analysis of the heart, kidney, liver, lung, and spleen showed no considerable alterations such as inflammation, necrotic sites, ischemia or hyperemia in the treated mice compared to the controls (Figure 3e). Blood chemistry further demonstrated no significant changes in alkaline phosphatase (ALP), aspartate aminotransferase (AST), or creatine kinase activity (Figure 3f), thus confirming the lack of adverse effects of Bola/IM on normal tissue. These findings suggest that Bola/IM has potent and selective anticancer activity against ovairan CSCs *in vivo* with no toxicity.

**Figure 3.**
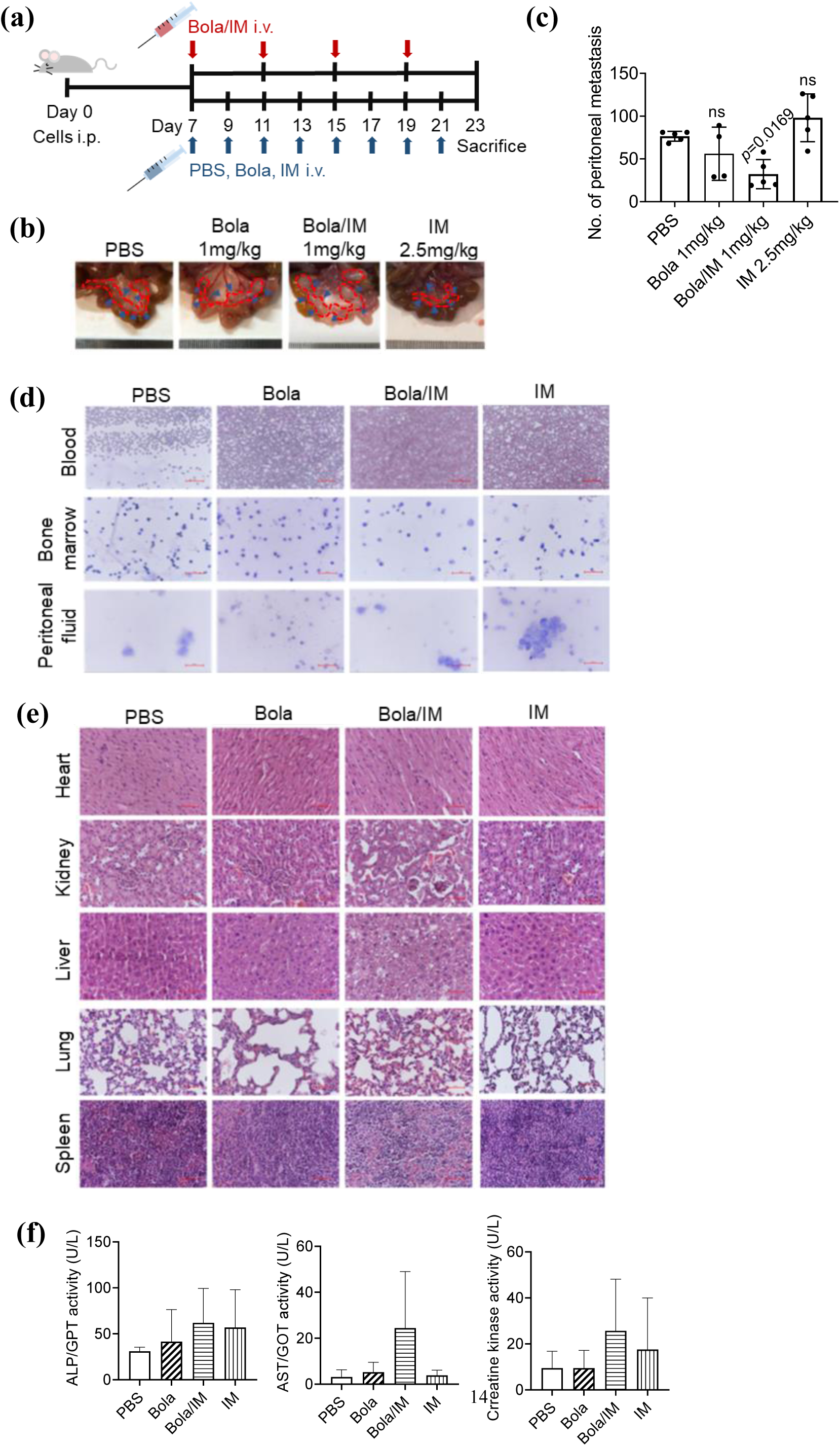
Therapeutic and chemosensitization effects of Bola/IM nanoformulation *in vivo*. (a) Timeline of treatments in tumor bearing mice (n=5). (b) Bola/IM nanoformulation inhibits peritoneal metastasis of HEYA8 CSCs at the dose of 1 mg/kg. Mesentery metastasis comparison. Arrows indicate large tumor nodes and (c) peritoneal metastasis number comparison after treatment. (d) Wright stained smears of blood, bone marrow, and peritoneal fluid. (e) Morphological characteristics of H&E stained organs and tissues following different treatments including Bola/IM nanoformulation. (f) Blood plasma biochemical markers evaluation. Experiments were repeated three times and data are shown as mean ± SD. Experiments were repeated three times and data are shown as mean ± SD. **, *P*<0.01 vs control. ***, *P*<0.001 vs control

### Bola/IM Achieves a High Tumor-Oriented Accumulation with Enhanced Drug Delivery Efficiency

To gain a better understanding of the promising antitumor activity exhibited by Bola/IM, we co-encapsulated the near-infrared fluorecent probe DiR within the Bola/IM nanoparticles (referred to as Bola/IM/DiR hereafter) to track their biodistribution *in vivo* in tumor-bearing xenograft mice. It should be mentioned here that we could not use DiR-labeled IM as control in our biodistribution study since it would not be representative of IM, the structural properties of the small molecule IM being largely overshadowed by those of the much larger DiR when conjugated. Our results showed that Bola/IM/DiR exhibited a much stronger and more localized fluorescence signal in tumor compared to the PBS and naked DiR controls at 8 h, with peak fluorescence observed at 48 h post intravenous injection (Figure 4a, b). This indicates that the bola-dendrimer nanoformulation prolonged the retention of loaded cargos in the circulating system, thereby facilitating their tumor-oriented accumulation *via* the EPR effect. Importantly, the DiR-labeled Bola/IM also penetrated deeper within the tumor tissues and spread further around the blood vessels compared to free DiR, as indicated in Figure 4c. This finding is particularly important for targeting CSCs most of which are deeply buried within the bulky tumor tissue.

**Figure 4.**
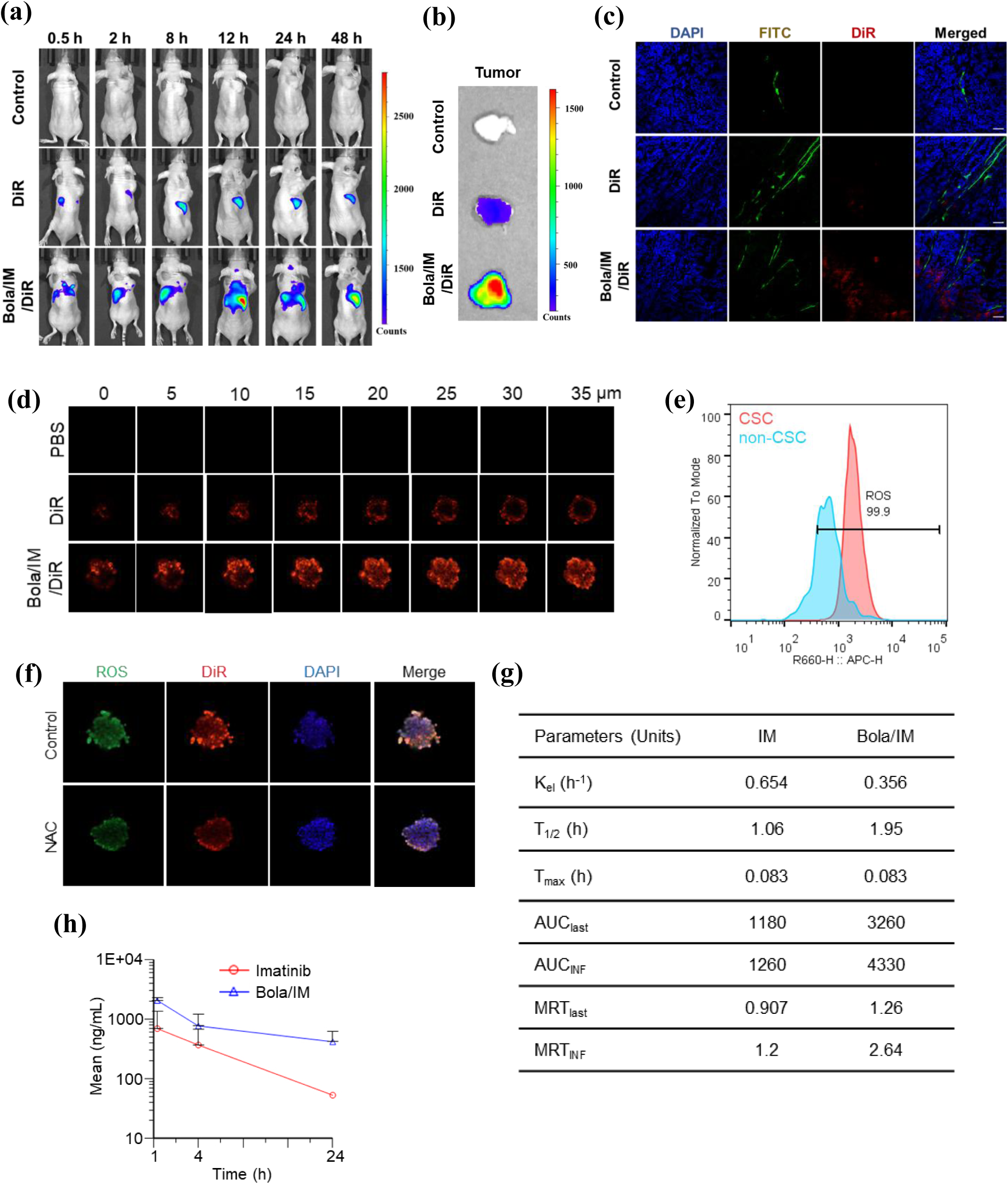
Bola/IM achieves a high tumor-oriented accumulation with enhanced drug delivery efficiency. (a) Nanoparticles incorporated with fluorescent dye were intravenously injected into mice with subcutaneous tumors, and imaged at different time points (0.5 h, 2 h, 4 h, 8 h, 12h, 24 h and 48 h). (b) Tumor were then harvested at 48 h after i.v. injection, the fluorescent signals of tumors were imaged. (c) Significantly improved accumulation and penetration within the tumor lesion. Red signal, DiR; green signal, FITC-tagged CD31 (staining tumor vessels) to show the excellent accumulation penetration of Bola/IM deep within the tumor lesion. (Scale bar: 50 μm.) (d) Penetration of Bola/IM/DiR within tumor spheroids. (e) APC-H histograms showing the ROS levels of CSC and non-CSC. (f) NAC inhibits ROS recruited Bola/IM/DiR accumulation in tumor spheroids. (g) Th1e6pharmacokinetic parameters of IM and of Bola/IM in mice following intravenous administration. (h) Plasma concentration-time profile of IM after administration of IM and Bola/IM to mice, respectively. Experiments were repeated three times and data are shown as mean ± SD.

We also used confocal microscopy to further examine the penetration of the DiR-encapsulated Bola/IM (Bola/IM/DiR) within tumor spheroids under nonadherent conditions, which most closely mimics malignant ascites in advanced/metastatic stages of ovarian carcinoma. Fluorescence imaging revealed significantly stronger and deeper penetration of Bola/IM/DiR within the spheroids compared to the controls of PBS and DiR alone controls (Figure 4d). These findings were consistent with our *in vivo* studies (Figure 4c) and further demonstrate the ability of Bola/IM to penetrate deep within tumor, where deep-seated CSCs can be effectively targeted.

Previously we have demonstrated that Bola-dendrimers possess favorable properties in response to ROS-rich conditions [25,26]. Cancer cells have been shown to have higher levels of ROS compared to normal tissue [30]. ROS signaling, by driving epithelial-mesenchymal transition, can induce cells to acquire stem-like properties [31]. In our study, we observed higher levels of ROS in HEYA8 CSCs compared to non-CSCs (Figure 4e). Interestingly, the fluorescent labeled Bola/IM (Bola/IM/DiR) concomitantly colocalized with regions of high ROS expression in CSCs and its levels decreased upon inhibition of ROS with NAC (Figure 4f; Figure S5). These findings confirm the higher uptake of Bola/IM in CSCs with high ROS levels and may also partly explain the effectiveness of Bola/IM against CSCs and not non-CSCs. Indeed, Bola/IM may be able to selectively target and eradicate CSCs in ovarian cancer by exploiting the elevated ROS levels in these cells.

Further investigation into the *in vivo* kinetic behavhior of Bola/IM revealed that Bola/IM had a 2.8-fold higher absorption (IM AUC_last_=1180, Bola/IM AUC_last_= 3260) and 50% lower elimination rate (IM K_el/h_=0.654, Bola/IM K_el/h_=0.356) compared to naked IM, suggesting a higher bioavailability and longer circulation retention of Bola/IM. Importantly Bola/IM reached the maximum concentration (T_max_) in the blood in a shorter amount of time and took longer to be reduced by half (T_1/2_) compared to naked IM, confirming the prolonged duration of Bola/IM in the body (Figure 4g). In addition, the mean residence time (MRT) of Bola/IM was 1.4-fold longer (Figure 4g), with a higher mean concentration at 24 h compared to naked IM (Figure 4h), indicating that less frequent treatment with Bola/IM at lower doses could lead to the desired therapeutic effects. These results demonstrate the promising tumor targeting ability of Bola/IM with enhanced drug delivery efficiency, further highlighting its potential as a promising therapeutic strategy for ovarian cancer.

### Chemosensitizing Effects of the Bola/IM Nanoformulation *In Vivo*

We previously found that c-Kit inhibition could reduce chemoresistance of ovarian CSCs. In this study, we established a patient-derived xenograft mouse model to examine whether Bola/IM could sensitize ovarian cancer to chemotherapeutics (Figure 5a). We found that combination of the first-line chemotherapeutic agent for ovarian cancer, cisplatin, with Bola/IM produced synergistic actions to suppress metastatic tumor growth (Figure 5b). We obtained similar findings using patient-derived organoids (Figure 5c). Immunochemical analysis showed a decrease in expression levels of two downstream interaction partners of c-Kit, HRP2 and β-catenin, in tumor tissues upon Bola/IM treatment (Figure 5d), which were further reduced upon combination of Bola/IM with cisplatin treatment (Figure 5d). Moreover, c-Kit expression is higher in patients who died of disease (DOD) compared to those alive with no evidence of disease (NED) (Figure S6). Collectively, these results confirm the effective anticancer activity of Bola/IM and the synergistic effect of combining it with cisplatin in treating ovarian cancer.

**Figure 5.**
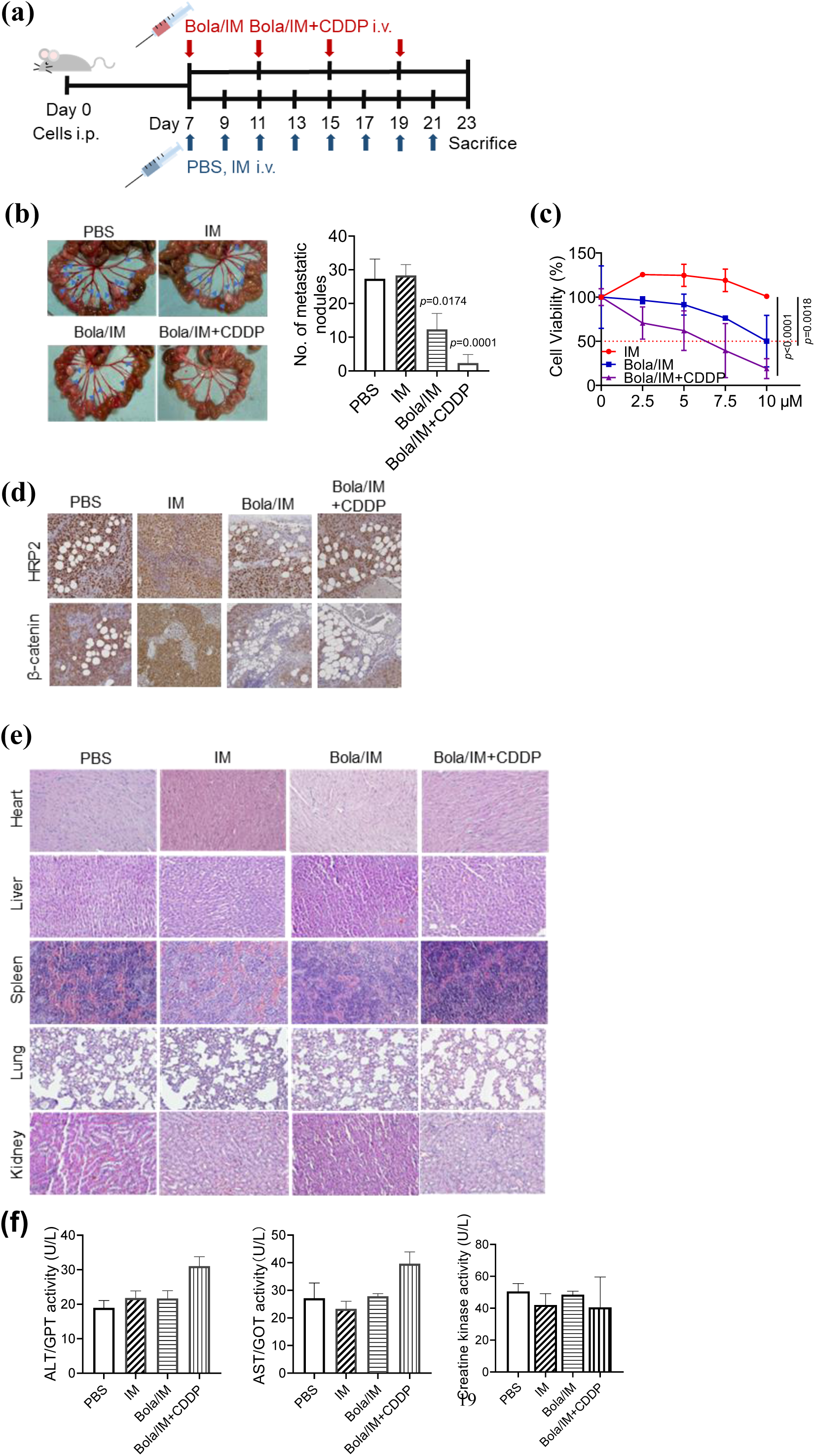
Bola/IM alone and in combination with CDDP decreases malignancy in patient-derived models without severe adverse effects. (a) Timeline of treatments in tumor bearing mice (n=5). (b) Cell viability of patient-derived organoids. (c) Mesentery comparison with arrows indicating large tumor nodes; the bar chart showing the number of peritoneal metastatic nodules according to treatment. (d) HRP2 and β-catenin expression levels were detected by immunohistochemistry in the tumor tissues. (e) Morphological characteristics of organs and tissues of mice treated with Bola/IM alone and in combination with CDDP. H&E stained organs and tissue comparison. (f) Blood plasma biochemical markers evaluation were measured and compared to control injected mice. Experiments were repeated three times and data are shown as mean ± SD. *, *P*<0.05 vs control. **, *P*<0.01 vs control. ***, *P*<0.001 vs control.

To further evaluate the potential toxicity of Bola/IM and its combination with cisplatin, we conducted histopathological analyses of major organs and blood biochemistry analysis in treated mice. Hematoxylin and eosin staining of the heart, liver, spleen, lungs, and kidneys showed no histological alterations (Figure 5e). Blood biochemical analysis revealed that neither Bola/IM nor its combination with cisplatin caused any significant increase in ALP, AST, or creatine kinase (Figure 5f), indicating the absence of damage to liver and kidney functions or any adverse effects on normal tissues. In addition, we observed no significant weight loss in the mice during the treatment (Figure S7). Altogether, these data confirm the safety and potential efficacy of Bola/IM as a nanoformulation for the treatment of ovarian cancer, addressing an urgent yet unmet medical need.

## DISCUSSION

Targeting CSCs, which are associated with drug resistance and recurrence in many tumors, has emerged as a promising treatment strategy for various cancers, including metastatic ovarian cancer. However, targeting CSCs has proven to be challenging due to their rarity, quiescent nature, and deep location within bulky tumors. Considering that c-Kit has been identified as a functional marker and a potential target for CSCs, and that the clinical anticancer drug imatinib (IM) targets c-Kit, we developed a nanodrug system based on the bola-amphiphilic dendrimer Bola to effectively deliver IM and enhance its anticancer activity on CSCs. We utilized *in vitro* and *in vivo* metastatic ovarian cancer models to evaluate the efficacy of this approach. Despite clinical trials of IM on patients with primary or recurrent ovarian cancer yielding unsatisfactory results, our nanodrug system holds promise for effectively targeting CSCs and improving IM treatment of ovarian cancer Dendrimers are nanosized molecules characterized by precise structure and cooperative multivalence. In this study, we demonstrated the effectiveness of a bola-amphiphilic dendrimer for the delivery of IM. This delivery system offers several advantages: i) high drug loading capacity, allowing for effective therapeutic potency, ii) small size (ca 30 nm), enabling passive tumor targeting and deep tumor penetration to reach the deep-seated CSCs; iii) higher uptake in CSCs with high levels of ROS, potentially enhancing the selective targeting of CSCs; iv) overcomes drug efflux and drug resistance; v) inhibits the β-catenin/HRP2 signaling pathway, which is implicated in CSC maintenance; and vi) effective anticancer activity, both as a standalone treatment or in combination with the clinical drugs paclitaxel and cisplatin. These findings highlight the successful implication of dendrimer nanotechnology in combating CSCs to overcome drug resistance and enhance the potency of anticancer drugs.

Previous studies have reported various nanoformulations of IM using poly(lactide-*co*-glycolide) or gold nanoparticles to increase its efficacy and reduce its cytotoxicity [32, 33]. However, none of these studies have described an optimal delivery system that effectively targets tumor tissue and exhibits a favorable biodistribution and pharmacokinetic profile. Such a delivery system is crucial for enabling the effective anticancer activity of IM against metastatic ovarian cancer while minimizing its side effects. In this study, we have demonstrated for the first time that Bola/IM rapidly accumulated at the tumor sites *via* the EPR effect and penetrated deep within the tumor. Moreover, Bola/IM effectively promoted uptake into ROS-rich cancer cells and inhibited drug efflux pumps. Importantly, while Bola/IM enhanced the antitumor activity of IM, it did not cause noticeable side effects or prominent toxicity. These findings align with our previous studies, where encapsulation of anticancer drugs such as paclitaxel, doxorubicin and rapamycin within amphiphilic dendrimers significantly reduces off-target toxicity [22–24]. In addition, the higher AUC and slower drug clearance rate observed in our study are attributed to faster accumulation at the tumor site and longer retention of Bola/IM in circulation. This suggests that lower and less frequent dosing of IM may suffice when encapsulated within Bola, thereby reducing its potential toxicity.

Our proteomic profiling study revealed HRP2 as a novel target of β-catenin that is inhibited by Bola/IM, which is a significant finding. We demonstrated for the first time the essential role of HRP2 and its interaction with β-catenin in the expansion of CSCs and development of drug resistance. HRP2 is frequently overexpressed in human hepatocellular carcinomas [34]. As a transcriptional activator, HRP2 contains crucial domains in its N-terminal region, which are involved in recognition of active chromatin, regulation of the transcriptional and pro-survival activity, and interactions with other proteins. HRP2 is also overexpressed in various other human cancers [35–37], and serves as an independent prognostic marker in intrahepatic cholangiocarcinoma [38]. Interestingly, blocking HRP2 has been shown to enhance the antitumor activities of targeted drugs and chemotherapy, suggesting that targeting HRP2 may overcome drug resistance [39, 40]. Accordingly in ovarian cancer, high HRP2 expression is associated with poor clinical outcomes. In CSCs, the interaction between HRP2 and β-catenin is crucial for self-renewal and tumorigenesis. Our analysis demonstrated that IM inhibited their expression and interaction at much higher doses when used alone compared to when encapsulated within the Bola/IM nanoformulation. The relatively low expression of HRP2/β-catenin in normal tissues further supports the potential of targeting HRP2 as a safe and promising approach for treating metastatic ovarian cancer.

## CONCLUSIONS

In this study, we demonstrate that the nanoformulation Bola/IM can empower the clinical anticancer drug IM in treating metastatic ovarian cancer by targeting cancer stem cells via β-catenin-HRP2 signaling axis. Previous clinical trials of IM as monotherapy or combination therapy have yielded unsatisfactory results for treating patients with primary or recurrent ovarian cancer [41–43]. Despite the initial response of most (75%) ovarian cancer patients to chemotherapy, nearly all eventually relapse due to the development of drug resistance, resulting in poor response or failure of chemotherapy. The high expression of drug transporters, such as P-glycoprotein and ABCG2, is a major cause of drug resistance in ovarian cancer. We demonstrate in this study that Bola/IM can decrease the expression of P-glycoprotein and ABCG2, leading to a coordinated decrease in drug efflux and an increase in apoptosis. Furthermore, when administered in combination with the clinical drugs cisplatin and paclitaxel, Bola/IM acts to sensitize CSCs to these chemotherapeutic drugs [7]. Our finding of a synergistic effect when combining Bola/IM and cisplatin in metastatic ovarian cancer suggests that other combination treatments with nanodrug formulations warrant further investigation. Such a strategy is expected to enhance the therapeutic effectiveness of targeted or immunotherapeutic drugs while reducing the morbidity associated with conventional cytotoxic chemotherapy.

## MATERIALS AND METHODS

### Experimental Animals

All animal studies were approved by Committee on the Use of Live Animals in Teaching and Research (The University of Hong Kong) and the Animals (Control of Experiments) Ordinance of Hong Kong.

### Chemicals

IM was obtained from MedChemExpress (Township, USA). Cisplatin and paclitaxel were from Calbiochem (San Diego, USA). Vinblastine was purchased from Sigma (St. Louis, USA). N-acetyl-L-cysteine (NAC) was purchased from Calbiochem (Rahway, USA). For the dendrimers, Bola was synthesized according to the previous work [25,26].

### Drug Loading Efficiency and Encapsulation Efficiency

The drug loading efficiency and encapsulation efficiency of the dendrimers were calculated according to the following equations:

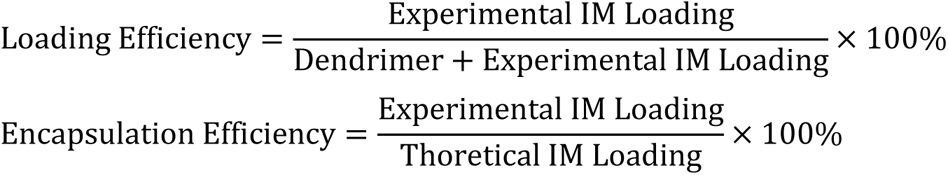

### Transmission Electron Microscopy (TEM)

Transmission electron microscopy was performed to characterize the size and morphology of Bola/IM complexes. 4.0 μL of solution containing Bola/IM was added onto a Formvar carbon-coated copper grid and air dried, followed by staining with 2.0% (w/v) uranyl acetate (Sigma-Aldrich, St. Louis, USA) and air dry before measurement. Isolated nanoparticles were fixed with 4.0% (w/v) neutral buffered formaldehyde and applied onto Formvar caron-coated copper grid for 20 min. The grids were post-fixed with 1.0% (w/v) glutaraldehyde (Sigma-Aldrich, St. Louis, USA), and then stained with 2.0% (w/v) uranyl acetate and allowed to air dry before measurement. TEM images were performed using JEOL 2100F analytical electron microscope (Tokyo, Japan) at an accelerating voltage of 200 kV.

### Nanoparticle Tracking Analysis (NTA)

Nanoformulations in liquid suspension were loaded into sample chamber of NanoBrookOmni (Brookhaven, USA) for particle size and zeta potential analysis.

### Cell Lines and Cell Culture

Human ovarian carcinoma cell line HEYA8 was a gift from Dr J Liu (MD Anderson Cancer Center, Houston, TX, USA) and SKOV3 were obtained from ATCC. Primary tumor samples were obtained from ovarian cancer patients with informed consent and approval by the University of Hong Kong Institutional Review Board. HEYA8 cells were maintained in Roswell Park Memorial Institute (RPMI) medium supplemented with 5% FBS (HyClone), and 100 units/mL penicillin/streptomycin at 37°C in a humidified incubator containing 5% CO_2_. SKOV3 were maintained in medium 199: MCDB105 (1:1) supplemented with 5% FBS, and 100 units/mL penicillin/streptomycin at 37°C in a humidified incubator containing 5% CO_2_. HEYA8 CSCs were used in modeling peritoneal metastasis animal model, and SKOV3 cells were used in *in vivo* biodistribution and penetration test.

### Cell Viability Assay

Cell viability was assessed using colorimetric MTT (3-(4,5-dimethylthiazol-2-yl)-2,5-diphenyltetrazolium bromide) assay following by the manufacturer’s instructions (Sigma). Briefly, 100 μL 10% MTT solution (5 mg/mL) was added to each well and plate was incubated for 4 h at 37°C. Medium was removed from each well and 100 μL DMSO was added. For tumor spheroids and organoids, samples were first collected and pelleted in tubes, 10% MTT solution were added and incubate for 4 h at 37°C. Then cell pellets were resuspended in 300 μL DMSO and transferred to 96-well plate. Analysis was performed at a wavelength of 570 nm by a microplate reader (Bio-Rad).

### Spheroid Formation Assay

Isolation and culture of HEYA8 and SKOV3 were performed as previously described. Briefly, 5000 cells/mL were plated in an ultralow attachment 100 mm culture plate in serum-free stem cell-selective medium. After 5-7 days, spheroid was generated and examined for subsequent assays.

### Reverse Transcription-Polymerase Chain Reaction (RT-PCR)

Total RNA was extracted with Trizol Reagent (Invitrogen) according to the manufacturer’s instruction. cDNA was synthesized using a First-strand cDNA synthesis kit (Invitrogen). PCR was performed with specific sequences of primers as following: Bmi1 Forward: 5’-ATG TGT GTG CTT TGT GGA G-3’, Reverse: 5’-AGT GGT CTG GTC TTG TGA AC-3’; Nanog Forward: 5’-AAG ACA AGG TCC CGG TCA AG-3’, Reverse: 5’-CCT AGT GGT CTG CTG TAT TAC-3’; Oct4 Forward: 5’-ATC CTG GGG GTT CTA TTT GG-3’, Reverse: 5’-TCT CCA GGT TGC CTC TCA CT-3’; β-actin Forward: 5’-TCA CCG AGG CCC CTC TGA ACC CTA-3’, Reverse: 5’-GGC AGT AAT CTC CTT CTG CAT CCT-3’.

### Western Blotting

Cells were collected in RIPA lysis buffer. Total proteins were separated by 7.5% separating SDS-PAGE gel and transferred to a nitrocellulose membrane. The proteins were blocked and incubated with primary antibodies at 4°C overnight followed by secondary antibodies after washes. Then protein signals were detected by Western Lightning Plus Enhanced Chemiluminescence (Perkin Elmer). Band intensities were calculated by densitometry using the ImageJ software. Antibodies were obtained as follows: P-glycoprotein (Abcam, ab202976, 1:1000), ABCG2 (CST, 4477, 1:1000), GAPDH (Abcam, ab8245, 1:5000), HRP2 (Invitrogen, XC3539332A, 1:1000), β-actin (Sigma, A5316, 1:5000), β-catenin (CST, 2698S, 1:5000), Goat anti-Rabbit IgG (H+L)-HRP conjugate (Bio-Rad, 1706515, 1:3000), Goat anti-Mouse IgG (H+L)-HRP conjugate (Bio-Rad, 1706516, 1:3000).

### Drug Efflux Test

Drug efflux test was performed using Multidrug Resistance Direct Dye Efflux Assay kit (Merck), according to manufacturer’s instruction. Briefly, cells were collected, disaggregated with 2 mM EDTA/PBS in a single-cell suspension with cold efflux buffer. Then, cells were incubated with DiOC_2_(3) solution. After that, cells were washed and distributed into 3 different tubes, containing vinblastine solution, DMSO, and cold efflux buffer. Tubes with DMSO and vinblastine were incubated at 37°C for 4 h and cold efflux buffer on ice all protected from light. Cells were then washed to stop the efflux reaction, stained with PI solution, and resuspended in PBS. Flow cytometry on BD FACSAria III (BD Biosciences) was performed, at excitation of 488 nm. FlowJo software was used for data analysis. Only live cells were included, and FITC signal was used as an indicator of DiOC_2_(3) efflux intensity.

### Immunoprecipitation-Mass Spectrometry

The nuclear extracts were prepared as described above. 500 μg nuclear extracts diluted in 500 μL IP lysis buffer was pre-cleared by incubating with 20 μL protein A/G agarose beads for 2 h at 4°C. Then the precleared nuclear proteins were mixed with 25 μL β-catenin primary antibody with Sepharose bead conjugated or 25 μL rabbit IgG control with Sepharose bead conjugated at 4°C overnight. The beads were washed by IP lysis buffer three times and pre-urea wash buffer (20 mM Tris pH 7.5, 150 mM NaCl). 50 μL urea elution buffer (8 M Urea, 25 mM ammonium bicarbonate) was added and incubated at 65°C for 15 min. The beads were pelleted by centrifugation, and the supernatant was transferred to a new 1.5 mL tube, followed by further trypsin digestion and peptide purification for MS analysis.

### Immunoprecipitation

The nuclear extracts were prepared as described above. 300 μg nuclear extracts diluted in 500 μL IP lysis buffer was pre-cleared by incubating with 20 μL protein A/G agarose beads (Santa Cruz) for 2 h at 4°C. Then the precleared nuclear proteins were mixed with primary antibodies overnight. 30 μL pre-washed protein A/G agarose plus-agarose beads were added to precipitate the antigen-antibody complexes at 4°C for 1 h. The beads were then washed by IP lysis buffer three times.

### Immunofluorescence Microscopy

Cells on coverslips were fixed with 4% paraformaldehyde/0.1% Triton, and blocked with 5% bovine serum albumin. Cells were incubated with primary HRP2 antibody (Invitrogen, 1:50) for 1 h, washed with PBS, and then incubated with secondary Cy3 antibody (ThermoFisher, 1:200) for 1 h. Signals were visualized using Carl Zeiss LSM 980 (Oberkochen, Germany).

### Small Interfering RNA Transfection

Non-specific and HRP2 siRNAs were purchased from Invitrogen. Cells were transfected with 20 nM siRNA using Lipofectamine 2000 following the manufacturer’s instructions (ThermoFisher).

### *In Vivo* Tumor Peritoneal Metastasis Assay

All mouse studies were performed in accordance with protocols approved by Committee on the Use of Live Animals in Teaching and Research (The University of Hong Kong) and the Animals (Control of Experiments) Ordinance of Hong Kong. 2×10^6^ cells were intraperitoneally (i.p.) injected into 6-8 weeks old female NOD/SCID mice. Body weights were measured every other day. For IM dosage valuation, mice were injected with PBS, IM 2.5 mg/kg or 5.0 mg/kg every two days for two weeks after one week of tumor cell injection. Mice were then sacrificed, and mesentery metastasis nodules were counted. For Bola/IM estimation, after one week following tumor cell injection, mice were randomly divided to receive PBS, empty Bola (1.0 mg/kg), Bola/IM (1.0 mg/kg), and naked IM (2.5 mg/kg). PBS, empty Bola and naked IM were treated every two days, Bola/IM were received at Day 7, 11, 15 and 19. PBS, empty Bola and naked IM were used as controls. For CDDP combination therapy, after one week following tumor cell injection, mice were randomly divided to receive PBS, naked IM (2.5 mg/kg), Bola/IM (1.0 mg/kg), and Bola/IM (1.0 mg/kg) + CDDP (5.0 mg/kg). PBS and naked IM were treated every two days, Bola/IM and CDDP combination were received at Day 7, 11, 15 and 19. PBS, empty Bola and naked IM were used as controls. At the end of experiments, mice were sacrificed and peritoneal metastases was counted. Tumor and organs (heart, lungs, liver, kidneys, spleen, brain, blood) were collected, fixed in formalin and embedded in paraffin for H&E and immunohistochemical staining. Blood plasma was collected for biochemical marker analysis using Infinity AST(GOT) kit, Infinity ALT(GPT) kit (ThermoFisher), Creatinine LiquiColor Test kit, CK-MB Liqui-UV Test kit, Direct Bilirubin LiquiColor Test kit (Stanbio) and Urea Assay kit (Abcam). Whole blood or peritoneal fluid were stained in Wright stain.

### *In Vivo* Biodistribution Study and Tumor Penetration

All procedures were approved by the Institutional Animal Care and Use Committee of China Pharmaceutical University and performed in accordance with the guidelines and policies. The approval number is “2023-09-009”. Female BALB/c nude mice (6-week-old) were purchased from Beijing Vital River Laboratory Animal Technology Co., Ltd. (Beijing, China). The mice were maintained in China Pharmaceutical University Laboratory Animal Center during the experiment, all mice were housed in specific pathogen-free conditions according to the current Chinese regulation of China Pharmaceutical University. 5.0×10^6^ SKOV3 cells were inoculated subcutaneously to the mice. After three weeks, the mice bearing SKOV3 xenograft tumors were randomly divided into 3 groups (n = 3/group), and then administrated with PBS (phosphate buffer saline), free DiR (40 μg/kg), and Bola/IM/DiR (4 mg/kg of Bola, 1 mg/kg of IM, 40 μg/kg of DiR) via tail vein of mice, respectively. The *in vivo* fluorescence images were recorded at 0.5, 2, 8, 12, 24, and 48 h after intravenous administration of Bola/IM/DiR. And the mice were visualized by using the small animal live optical imaging system (PerkinElmer, IVIS@ spectrum, USA). During imaging, mice were anesthetized with isoflurane gas, a 740 nm pulsed laser diode was used to excite the DiR molecules, emission wavelength signal at 770 nm was collected. Furthermore, Mice were sacrificed after 48 h, the excised organs (hearts, livers, spleens, lungs, kidneys) and tumors were collected for imaging. In the last, the tumor tissue was used to make frozen sections and immunohistochemical tests, blood vessels were labelled with FITC-labeled CD31, and nuclei with DAPI, sections were then photographed using laser scanning confocal microscope (ZEISS, LSM880, Germany), to detect the penetration of DiR-labelled Bola/IM in tumor lesion.

### Drug Penetration in Tumor Spheroids

The penetration ability of DiR-labelled Bola/IM in multicellular tumor spheroids was studied using Carl Zeiss LSM 980 (Oberkochen, Germany). Cells were seeded in low-attachment culture dish containing serum free medium allowing 14 days to form tumor spheres. 1.0 mg/mL Bola/IM/DiR were added into culture medium for 5 h, washed with PBS and fixed with 4% formaldehyde at 4℃ for 20 min. Spheroids were then resuspended in 60% glycerol after washing with PBS twice for z-stack imaging. The images were captured from the lower position top up to the center of spheroids every 5 μm at 10× magnification.

### Flow Cytometry

Bola/IM/DiR incubated CSCs were treated with NAC for 24 h, spheroids were collected and resuspended with 100 μL of 1× ROS Assay Stain (Invitrogen) for 60 min. Then spheroids were trypsinized to single cells for flow cytometry analysis (Agilent NovoCyte Quanteon).

### Pharmacokinetics Analysis

The bioanalysis condition was specified as follows: LC system (Shimadzu LC-40D system), MS system (Sciex QTRAP 6500+), column (Agilent Eclipse Plus C18, 18 μm, 2.1×50 mm), column temperature 40°C, flow rate 0.2 mL/min, electrospray ionization (positive). The mobile phase consisted of 0.1% formic acid in ultrapure water (mobile phase A) and 0.1% formic acid in acetonitrile (mobile phase B). The following gradient profile was used: (1) 1 min, 5% B; (2) 2.5 min, 70% B; (3) 6.5 min, 70% B; (4) 7.5 min, 5% B; (5) 8.0 min, 5% B. IM was extracted from serum by methanol. The samples were vortexed, centrifuged and the supernatant was then injected to LC-MS/MS. The calibration curves and quality control samples were prepared in blank mouse serum and treated following the same procedure as the samples.

### Immunohistochemistry Staining

Slides with tissue sections were rehydrated. Endogenous peroxidase activity was inhibited by incubation in 3% hydrogen peroxide. Antigen retrieval was performed with heating slides at 90°C in a citrate buffer (pH 6.0) and then cooling slides at a room temperature. Slides were then incubated with primary antibody solution in a blocking buffer overnight at 4°C, washed with TBS-T, and incubated with the biotin-conjugated secondary antibody. Slides were then incubated with streptavidin and with DAB solution and counterstained in hematoxylin (Biocare medical).

### Statistical Analysis

Each experiment was repeated at least 3 times. Normality of the sample distribution was checked with Shapiro-Wilk test. *t*-test was used for parametric samples, and Wilcoxon test was used for non-parametric samples with 2 groups. One-way ANOVA was used for multiple comparisons. Two-way ANOVA was used for multiple groups comparison with 2 or more categorial variables. *p*<0.05 was considered as statistically significant.

### Data Availability Statement

The data that support the findings of this study are available from the corresponding authors upon reasonable request.

## Acknowledgements

This work was supported by the Health and Medical Research Fund 06173496 (A.S.T.W.) and the funding support from “Laboratory for Synthetic Chemistry and Chemical Biology” under the Health@InnoHK Program launched by Innovation and Technology Commission, HKSAR.

## Notes

The authors declare that they have no competing interests.

## Author Contributions

Z.S. and M.A. contributed equally to this work. Z.S., M.A. and M.Z. performed cellular experiments. Z.S., M.A., M.Z. and S.L. performed animal experiments. Z.S. and M.A. contributed to conceptualization and methodology. Z.S. wrote the original draft. Z.S., X.L., P.L., and A.W. reviewed and edited the manuscript. Z.B., B.L., F.M., J.C., T.R. and Y.L. performed nanoparticle synthesis and drug encapsulation. K.C., P.I., and H.C. provided clinical samples. X.L., L.P., and A.W. supervised the project.

