## Supplementary Figures for "Bola-amphiphilic dendrimer empowers imatinib to target metastatic ovarian cancer stem cells *via* β-catenin-HRP2 signaling axis"

### Supporting figures

**a**

| Formulation | Dendrimer/Imatinib (W/W) | Drug Loading (%) | Encapsulation Efficiency (%) |
|---|---|---|---|
| | 16/2 | 10.7 | 96.9 |
| Bola/IM | 16/4 | 19.5 | 99.2 |
| | 16/8 | 33.2 | 99.7 |

**b**

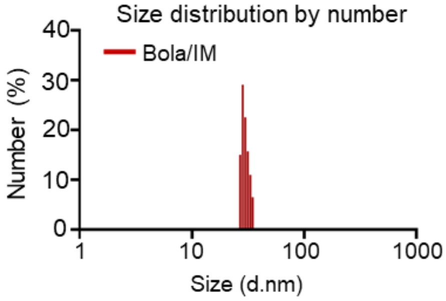


**c**

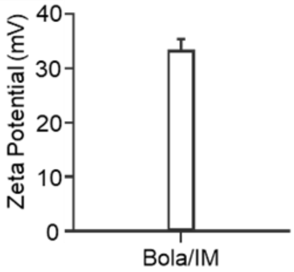


**d**

| $IC_{50}$, μM | Imatinib (IM) | Bola/IM 16/4 | Bola |
|---|---|---|---|
| HEYA8 non-CSCs | 45.2 | 18.8 | 76.2 |
| HEYA8 CSCs | 7.5 | 1.6 | 154 |
| SKOV3 non-CSCs | >100 | 96.2 | 230 |
| SKOV3 CSCs | >100 | 5.7 | 29.6 |
| OSE | >100 | 38.6 | 52.1 |

Figure S1. Nanoformulation of IM using the bola-amphiphilic dendrimer (Bola/IM) specifically targeted HEYA8 CSCs. (a) The drug loading content and encapsulation efficiency. (b) size distribution and (c) zeta potential of Bola/IM nanoformulation. (d) *In vitro* cytotoxicity $IC_{50}$ (μM) values of IM, Bola and Bola/IM 16/4 towards non-CSCs and CSCs. Experiments were repeated three times.

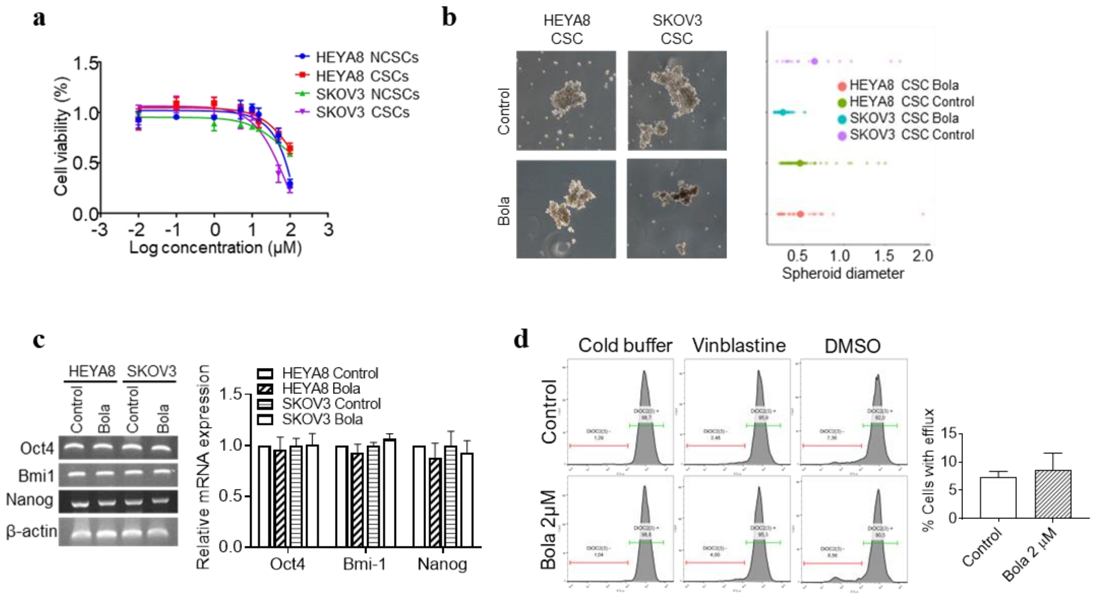


Figure S2. Bola empty nanoparticle has no anti-CSC effects and does not reduce drug efflux. (a) *In vitro* cytotoxicity $IC_{50}$ (μM) value of Bola towards non-CSCs and CSCs. (b) CSCs were treated with Bola for 7 days. The diameter of tumor spheres was measured. (c) No changes in stemness markers were identified by RT-PCR. β-actin was used as a control. (d) No influences on drug efflux in HEYA8 CSCs treated with Bola. Experiments were repeated three times and data are shown as mean ± SD.

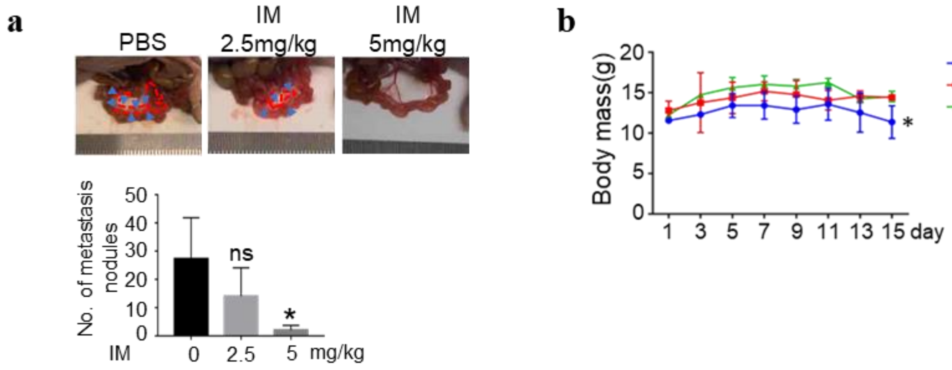


Figure S3. Naked IM inhibits peritoneal metastasis at the dose of 5 mg/kg. (a) Mesentery comparison. Arrows indicate large tumor nodes and peritoneal metastasis number comparison after treatment. (b) Mice body weights were monitored every 2 days. Experiments were repeated three times and data are shown as mean ± SD. *, $P <$ 0.05 vs control.

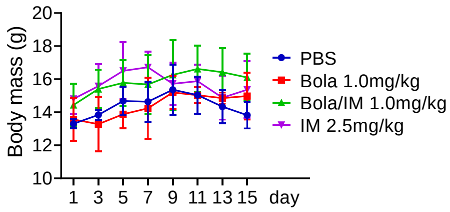


Figure S4. No body weights loss was caused upon Bola/IM treatments. Mice body weights were monitored every 2 days. Experiments were repeated three times and data are shown as mean ± SD.

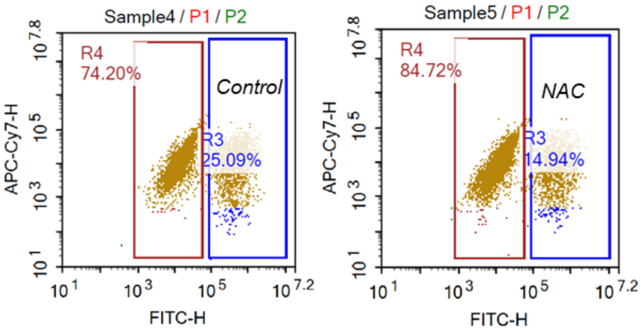


Figure S5. NAC inhibits ROS recruited Bola/IM/DiR accumulation in tumor spheroids. FITC-H histograms showing the ROS levels of CSCs incubated with/without NAC.

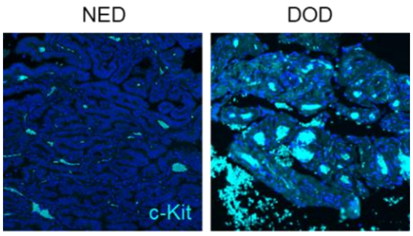


Figure S6. High c-Kit (cyan) expression in patient tissues indicates poor outcome. NED: alive with no evidence of disease; DOD: died of disease. c-Kit signal (cyan) was visualized under confocal microscopy.

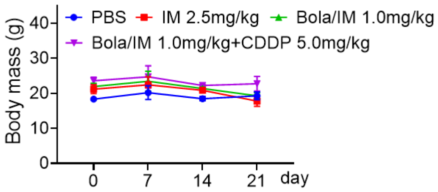


Figure S7. No body weights loss was caused upon Bola/IM nanoformulation and combination treatments. Mice body weights were monitored every 7 days. Experiments were repeated three times and data are shown as mean ± SD.
